# BCLncRDB: A comprehensive database of LncRNAs associated with breast cancer

**DOI:** 10.1101/2022.12.05.519223

**Authors:** Swapnil Kumar, Avantika Agarwal, Vaibhav Vindal

**Affiliations:** Department of Biotechnology & Bioinformatics, School of Life Sciences, University of Hyderabad, Gachibowli, Hyderabad, 500046, India

## Abstract

**Motivation:** Breast cancer, the most common cancer in women, is characterized by high morbidity and mortality worldwide. Recent evidence has shown that long non-coding RNAs (lncRNAs) play a crucial role in the development and progression of breast cancer. Despite this, no database exists primarily for lncRNAs associated with only breast cancer.

**Results:** We developed BCLncRDB, a manually curated, comprehensive database of lncRNAs associated with breast cancer. For this, we collected, processed, and analyzed data on breast cancer-associated lncRNAs from different sources, including published literature and TCGA. Currently, our database contains 5,279 unique breast cancer-lncRNA associations. It has the following features: (I) Differentially expressed and methylated lncRNAs, (II) Stage and subtype-specific lncRNAs, and (III) Drugs, Subcellular localization, Sequence, and Chromosome information. Thus, the BCLncRDB provides a dedicated platform for exploring breast cancer-related lncRNAs to advance and support the ongoing research on this disease.

**Availability and implementation:** The database BCLncRDB is publicly available for use at http://sls.uohyd.ac.in/new/bclncrdb.

## 1. Introduction

Long non-coding RNAs (lncRNAs) are a class of non-coding RNA (ncRNA) molecules with a length that ranges from 200 base pairs (bp) to 100 kilobases (KB) [Kung et al., 2013; Li et al., 2013]. Initially, this class of ncRNAs is thought to be noisy without any biological functions. However, recent studies have indicated that lncRNAs are involved in various cellular physiological regulations such as polypeptide encoding, epigenetic regulation, transcriptional regulation, post-transcriptional regulation, and signal transducer [Jin et al., 2021]. For example, lncRNA MEG3 suppresses the regulatory activity of the *MDM2* gene, a negative regulator of the *P53* gene, via phosphorylation resulting in tumor suppression by *P53*. Similarly, lncRNA LOC572558 acts as a negative regulator of *MDM2*, leading to tumor suppression by *P53* [Zhu et al., 2016]. Moreover, experimental evidence suggests the prominent roles of lncRNAs in the development and tumorigenesis of various cancers [Kopp et al., 2018; Pang et al., 2019; Sun et al., 2015; Chen et al., 2019].

Breast cancer is a highly heterogeneous disease with multiple subtypes and a major cause of concern for females at risk around the globe [Sung et al., 2021]. Currently, available treatment regimes are insufficient to tackle the problem that arises due to metastasized tumors, disease recurrence, limited subtype-specific treatment options, and resistance development. Thus, there is an urgent need for an in-depth understanding and exploration of other regulatory molecules, such as lncRNAs, known for their roles and potentials associated with breast cancer. This, in turn, may lead to the identification and development of novel, more effective, and subtype-specific therapeutic and diagnostic candidates.

Many databases on cancer-related lncRNAs have been developed to store a wide range of information available for cancer-related lncRNAs. For example, the Lnc2cancer database has information on lncRNAs and human cancer associations. All the lncRNAs-cancer association information stored at the Lnc2cancer is manually curated from the previously published literature [Ning et al., 2016]. Another database named LncRNADisease provides information on lncRNA-disease associations across various diseases, including cancers. These relationships have mainly been identified through *in vitro* experiments or computational predictions [Chen et al., 2012]. Next, the LnCaNet provides a resource for a comprehensive co-expression network between lncRNAs and non-neighboring cancer genes [Liu et al., 2016]. Furthermore, Lnc2catlas is a database that provides quantitative associations between lncRNAs and various cancers [Ren et al., 2018]. Although these databases are useful in studying and researching cancer-related lncRNAs, there is a lack of dedicated databases or web resources on lncRNAs associated with only breast cancer.

To this end, we developed a manually curated comprehensive database of breast cancer-associated lncRNAs, named BCLncRDB. For creating the database, firstly, we collected various information on lncRNAs associated with breast cancer, including expression and methylation patterns, targets, effects of drugs on lncRNA expression, sub-cellular localization of lncRNAs, sequence, and chromosomal location from previously published literature on lncRNAs in breast cancer, The Cancer Genome Atlas (TCGA) [Weinstein et al., 2013], Gene Expression Omnibus (GEO) [Barret at al., 2012], and various other sources including Ensembl database [Hubbard et al., 2002]. Secondly, this information on lncRNAs was stored with the help of MySQL, while the front end of the database was constructed with the use of HTML and PHP. We envision that BCLncRDB will be a productive resource designed explicitly for lncRNAs associated with breast cancer to accelerate future research in this area.

## 2. Results

### Database Content

More than 3000 published research articles on lncRNAs associated with breast cancer were retrieved from the PubMed database. These articles were systematically reviewed and curated, as discussed in the Materials and methods section. Only 936 articles were considered further for extracting information on lncRNAs, including 682 unique lncRNAs associated with breast cancer. After completing this process, data retrieved from these articles included the lncRNA name, Ensembl ID, chromosome information, expression patterns (upregulated, downregulated), experimental techniques (e.g., qRT-PCR), experimentally validated targets of lncRNAs, experimental samples (tissues, cell lines, etc.), subtype information of breast cancer (e.g., Her-2, Basal, Luminal A (LumA), Luminal B (LumB), and Normal-like (NormL)), drug information on lncRNA expression, subcellular localization of lncRNAs, and associated literature (PubMed ID, year of publication, and the title of the paper).

Moreover, publicly available microarray and RNAseq expression profile data of lncRNAs were also analyzed to identify differentially expressed lncRNAs in breast tumor tissues. Four GEO Datasets (GSE60689, GSE64790, GSE113851, and GSE119233) were found useful after filtering based on criteria as discussed in the Materials and methods section. Thus, we got 2,944 lncRNA-breast cancer associations from these GEO data, including 2,905 novel lncRNAs. The detailed information of GEO Datasets and differentially expressed lncRNAs retrieved by analyzing them has been described in Table 1.

**Table 1.** Distribution of samples and differentially expressed lncRNAs in different GEO datasets.

| GEO Accession | No. of Samples |  | LncRNAs |  |
| --- | --- | --- | --- | --- |
|  | Tumor | Normal | Downregulated | Upregulated |
| GSE64790 | 3 | 3 | 218 | 134 |
| GSE60689 | 2 | 2 | 420 | 300 |
| GSE113851 | 44 | 13 | 77 | 36 |
| GSE119233 | 20 | 10 | 702 | 1227 |

From the gene expression data of TCGA, we identified a total of 734 lncRNAs associated with breast cancer, out of which 720 lncRNAs were unique. In the case of the expression data, GEO, and TCGA, the association of lncRNAs with breast cancer has been determined based on the differential expression of these lncRNAs in tumor tissues as compared to adjacent normal tissues of the breast.

For stage-specific and subtype-specific lncRNAs, we collected the samples (Tumor and Normal Tissues) from TCGA as discussed in the Materials and methods section. Stage I, II, III, and IV contained 179, 607, 241, and 19 tumor samples, respectively, while subtypes *viz*. Lum A, Lum B, Her-2, Basal, and NormL contained 559, 201, 81, 187, and 40 tumor samples, respectively. Further, 112 adjacent normal tissue samples were used to identify the stage and subtype-specific lncRNAs (Table 2). The Limma R/Bioconductor package [Ritchie et al., 2015] was used to determine the differentially expressed stage-specific and subtype-specific lncRNAs. The adjP-value < 0.05 and |log2fold change| ≥ 2 were used as the criteria to identify upregulated and downregulated stage and subtype-specific lncRNAs. Thus, breast cancer stages *viz*. I, II, III, and IV had 193, 215, 213, and 252 differentially expressed lncRNAs, respectively, whereas subtypes *viz*. Lum A, Lum B, Her-2, Basal, and NormL had 228, 407, 405, 387, and 61 differentially expressed lncRNAs, respectively (Table 2).

**Table 2.**
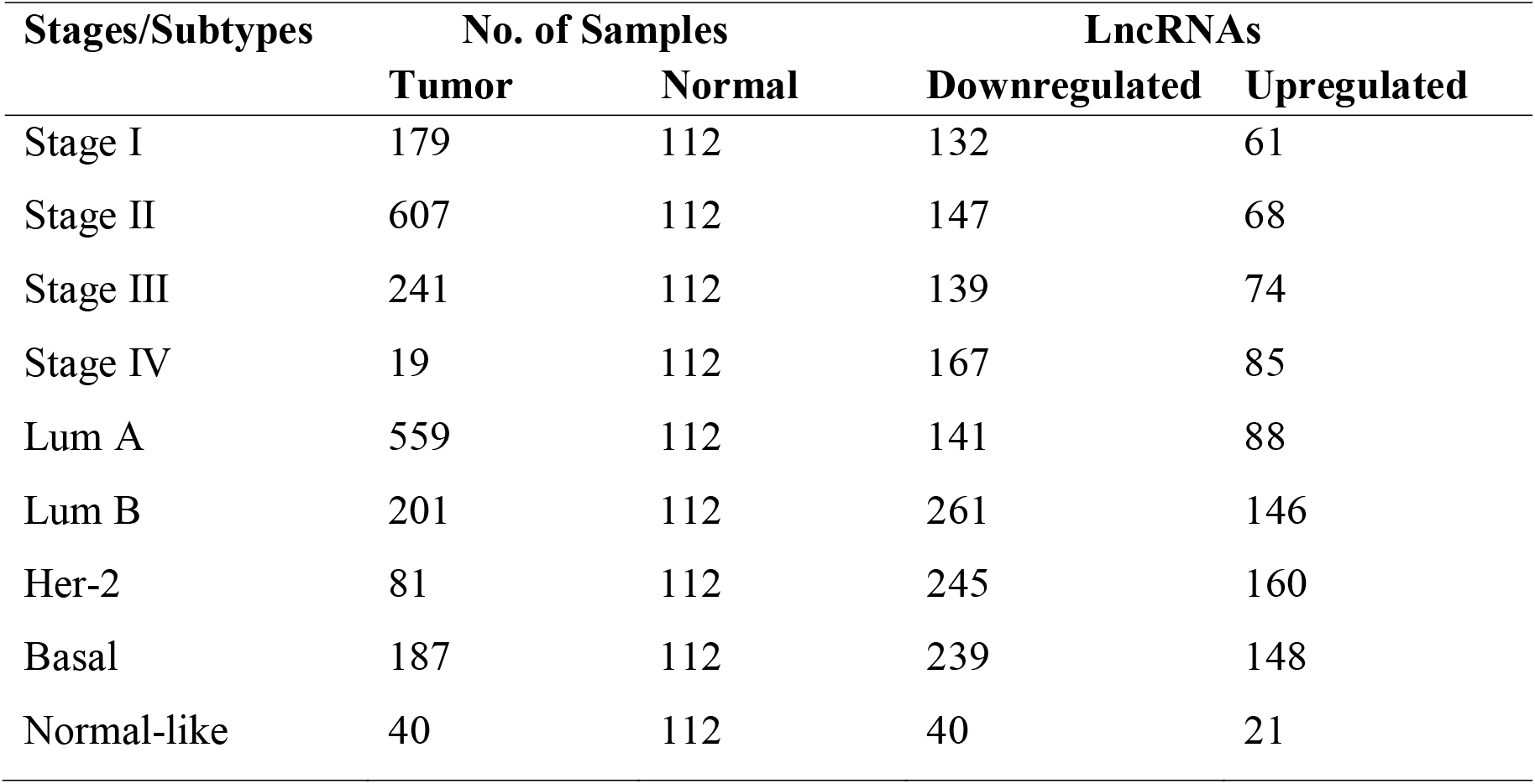
Distribution of samples and differentially expressed lncRNAs in TCGA data for different stages and subtypes of breast cancer.

From the above data, we got 12 lncRNAs which were present in all subtypes (Lum A, Lum B, Her-2, Basal, and NormL), and their expression pattern was upregulated; on the other hand, 27 lncRNAs were present in all subtypes (Lum A, Lum B, Her-2, Basal, and NormL) with expression pattern as downregulated. Further, 42 lncRNAs were present in all stages (I, II, III, and IV), having expression patterns as upregulated. In comparison, 101 lncRNAs were present in all stages (I, II, III, and IV), with expression patterns downregulated.

There were 9, 8, 14, and 63 lncRNAs specific to stages I, II, III, and IV, respectively, while 10, 66, 79, and 158 lncRNAs specific to subtype Lum A, Lum B, Her-2, and Basal, respectively. Detailed information on lncRNAs for stage-specific and subtype-specific is given in Table 3.

**Table 3.** Distribution of stage-specific and subtype-specific lncRNAs.

| Stage/Subtype | LncRNAs<sup>#</sup> |  |
| --- | --- | --- |
|  | Downregulated | Upregulated |
| Stage I | 6 | 3 |
| Stage II | 4 | 4 |
| Stage III | 8 | 6 |
| Stage IV | 33 | 30 |
| Lum A | 1 | 9 |
| Lum B | 39 | 27 |
| Her-2 | 25 | 53 |
| Basal | 85 | 73 |

No lncRNA was common among all three sources, i.e., literature, TCGA datasets, and GEO datasets. Among all downregulated lncRNAs, 187 lncRNAs from the literature, 401 lncRNAs from the TCGA dataset, and 1,292 lncRNAs from the GEO datasets. Similarly, among all upregulated lncRNAs, there were 487 lncRNAs from the literature, 306 lncRNAs from the TCGA dataset, and 1,650 lncRNAs from the GEO datasets.

When we compared the content of our database, i.e., BCLncRDB, with other publicly available databases on lncRNAs associated with diseases including breast cancer such as LncRNADisease, Lnc2Cancer, lnCaNet, and Lnc2catlas, it was found that the BCLncRDB hosts a wide variety of information on lncRNAs associated with breast cancer and this information are not available in any of these prior-available databases or web-resources. For example, information such as drug, methylation, subtypes, and stages of breast cancer and associated differential lncRNAs are unique features of our database. The total number of lncRNAs associated with breast cancer, as reported in our database as a single public platform, is also comparatively higher than the number of lncRNAs associated with breast cancer as available in any other databases or web resources. A detailed comparison of all available information on the disease-associated lncRNAs across different databases is available in Table 4.

**Table 4.**
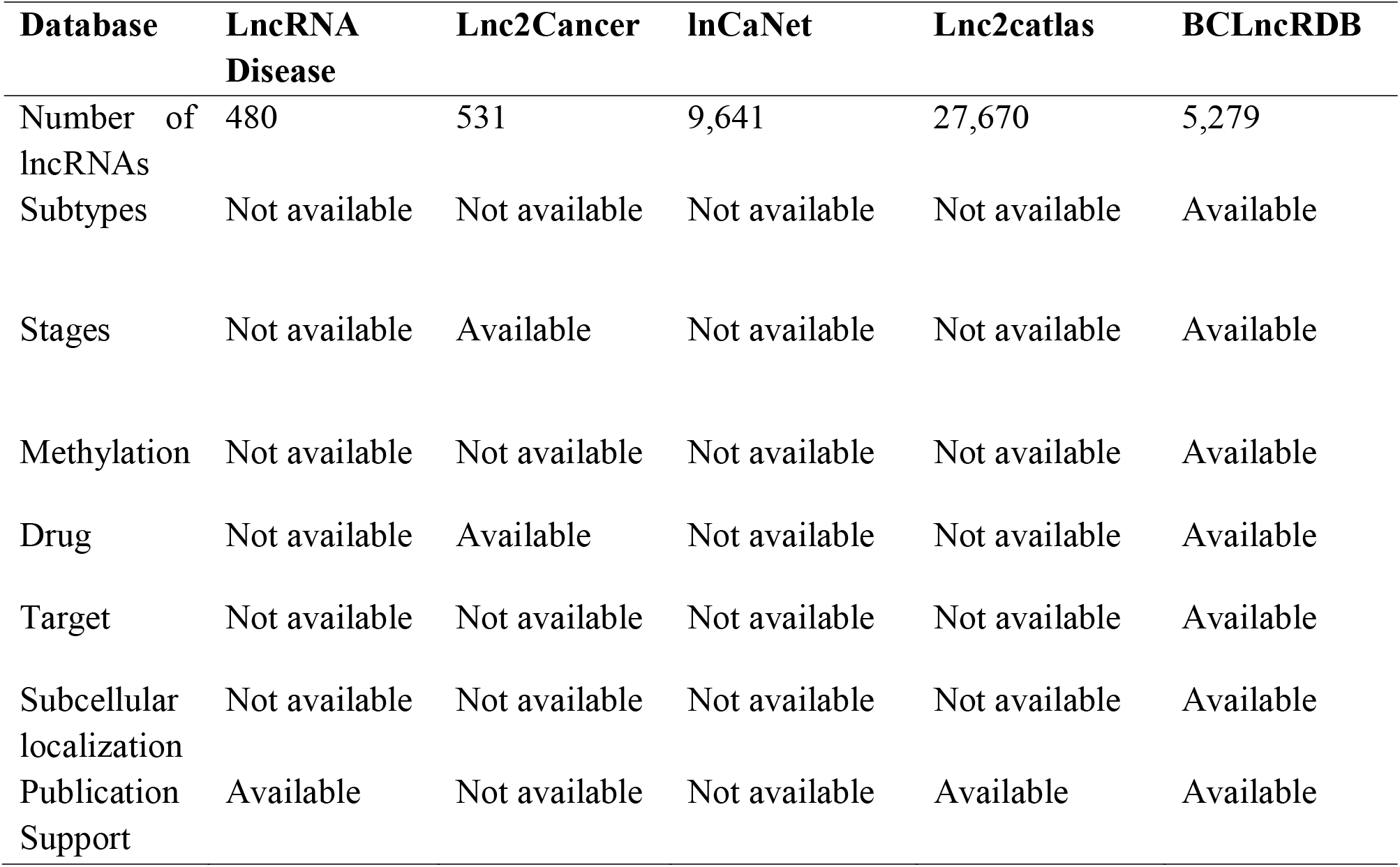
Comparison among BCLncRDB and five other lncRNA-associated cancer databases.

### Database Interface

The BCLncRDB, freely available at http://sls.uohyd.ac.in/new/bclncrdb, provides several search functionalities on the navigation bar for end users to retrieve useful information on lncRNAs available therein. By clicking on any search option, users can enter the lncRNA name or Ensembl ID of interest in the search box to retrieve various information about that particular lncRNA. For a specific lncRNA name or Ensembl ID, the following details in the general search option will be displayed (Fig. 1A and 1B): lncRNA name, Ensembl ID, chromosome information, expression patterns of lncRNAs, experimental techniques, experimental samples, stage, subtype information of breast cancer, PubMed ID, year of publication, and title of the paper. The lncRNA-target search option will display the following details (Fig. 1C and 1D): lncRNA name, Ensembl ID, target, regulatory direction, experimental method for lncRNA target, expression patterns, experimental method for lncRNA expression, lncRNA position, the title of the paper, and Pubmed ID. Next, the methylation search option will display the following information (Fig. 1E and 1F): lncRNA name, Ensembl ID, methylation pattern, and source. Further, the drug-target search option will provide the following information (Fig. 1G and 1H): lncRNA name, Ensembl ID, expression pattern, method, target gene, pathway, drug ID, drug name, drug method, drug response, experimental samples, the title of the paper, and paper’s Pubmed ID.

**Fig. 1.**
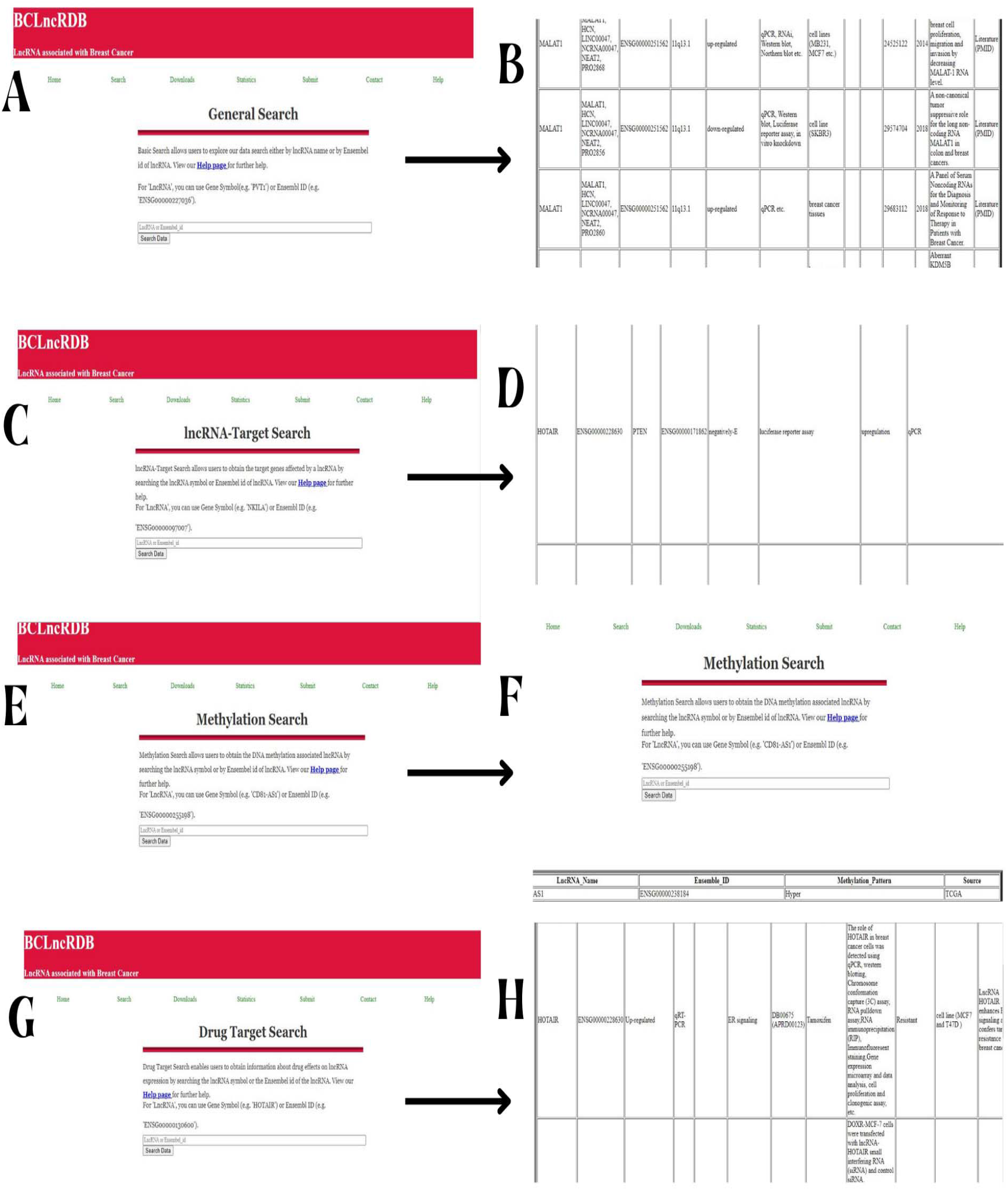
Screenshot of the search pages

Users can also retrieve the complete data provided in the database from the Download page available at the database interface. On the Download page, users can download the data in XLS format. Further, a help page provides detailed information for any user to ensure easy access to the data as a first-time visitor to the database.

### Data Statistics

All lncRNAs associated with breast cancer were collected from various sources, *viz*., GEO, lncLocator [Lin et al., 2021], Published Literature, and TCGA. Among the total lncRNAs, 5279 lncRNAs were identified as unique lncRNAs associated with breast cancer. Figure 2 shows the detailed distribution of these lncRNAs across different sources as a histogram.

**Fig. 2.**
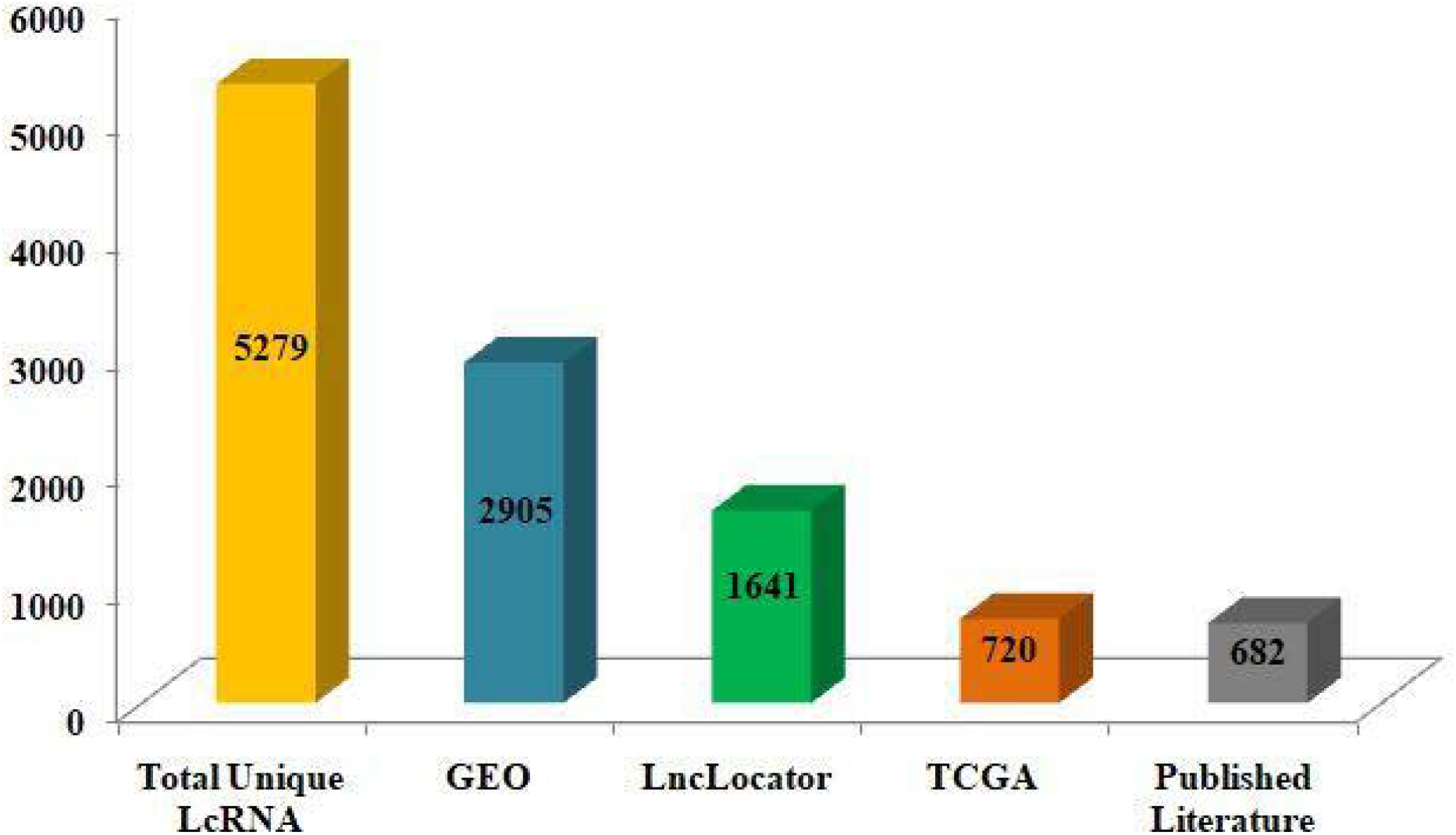
LncRNA distribution across the different sources

Further, a total of 25 lncRNAs were found differentially methylated in the case of breast cancer from TCGA data (Fig. 3). Of these 25 lncRNAs, 15 were hypo-methylated, while ten were hyper-methylated. Moreover, target information of 132 lncRNAs associated with breast cancer was retrieved from published research articles. Also, 20 unique lncRNAs related to breast cancer were found to have drug information based on lncRNA expression as obtained from published literature. A total of 1,653 unique lncRNAs have information on subcellular localization as collected from published literature and the lncLocator database (Fig. 3). The BCLncRDB also hosts chromosome and sequence information of 1,419 and 755 unique lncRNAs, respectively.

**Fig. 3.**
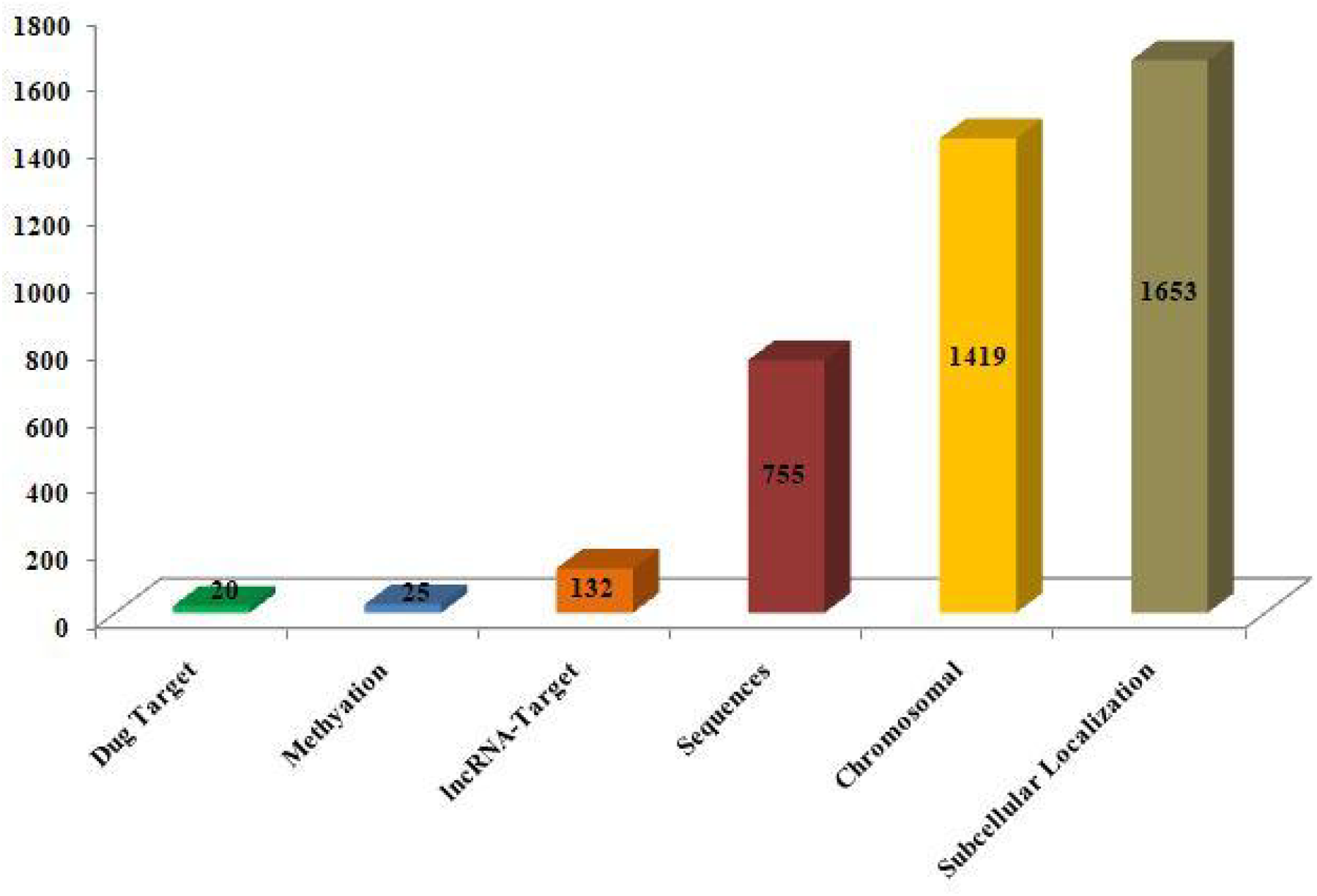
Distribution of lncRNAs with Methylation, Drug-Target, lncRNA-Target, Chromosomal, and subcellular localization information.

## 3. Discussion

The BCLncRDB (http://sls.uohyd.ac.in/new/bclncrdb/) is the first dedicated database on lncRNAs associated with breast cancer that stores a large number of information such as lncRNA name, Ensembl ID, breast cancer subtype, breast cancer stage, expression pattern, methylation pattern, chromosomal location, targets, pathway, drug information, sequence, sub-cellular localization, Pubmed ID, and experimental techniques. Our breast cancer-associated lncRNA database can be useful for researchers working on breast cancer in various ways: (i) researchers can access breast cancer-associated lncRNAs at stage and subtype levels, along with expression and methylation patterns that can be utilized for exploring stage and subtype-specific biomarkers and therapeutic candidates; (ii) they can seek drugs, resistance, and targets information to infer more efficient and new drug targets; and (iii) users can download the lncRNAs data such as targets, methylation, drugs, sequence and subcellular localization for other related studies of their interest. Further, as the high-throughput sequencing costs have decreased and technologies have advanced, more tumor tissues paired with adjacent normal or normal tissue samples will be sequenced in the near future. We will continue to make enhancements to the database content, such as the addition of extra information on lncRNAs associated with breast cancer, including newly reported lncRNAs in breast cancer, simple nucleotide variation (SNV), and copy number variation (CNV), along with missing information on existing lncRNA entries of the database.

In summary, we developed a unique and comprehensive database of lncRNAs associated with breast cancer named “BCLncRDB.” It not only provides the experimentally validated data but also provides the information from the TCGA dataset, GEO datasets, and Ensembl database. The development of BCLncRDB aims to provide a more orientated, curated database of lncRNAs associated with breast cancer that hosts information on subtypes, stages, gene expression, methylation, sequences, chromosomal location, targets, drugs, and many others. Several studies suggest that the deregulation of lncRNA has a vital role in the development and progression of breast cancer. Thus, we believe that this comprehensive and expandable resource of lncRNAs associated with breast cancer will facilitate scientists with an extended platform to accelerate breast cancer research to identify more effective and specific therapeutics for the disease.

## 4. Materials & methods

Various information on lncRNAs associated with human breast cancer was collected to develop the database. This information was collected not only from published literature but also from publicly available data (like RNA-seq, microarray, methylation, etc.) on lncRNAs available at various databases such as Gene Expression Omnibus (GEO) of NCBI and The Cancer Genome Atlas (TCGA). Figure 4 illustrates the schematic schema of database construction.

**Fig. 4.**
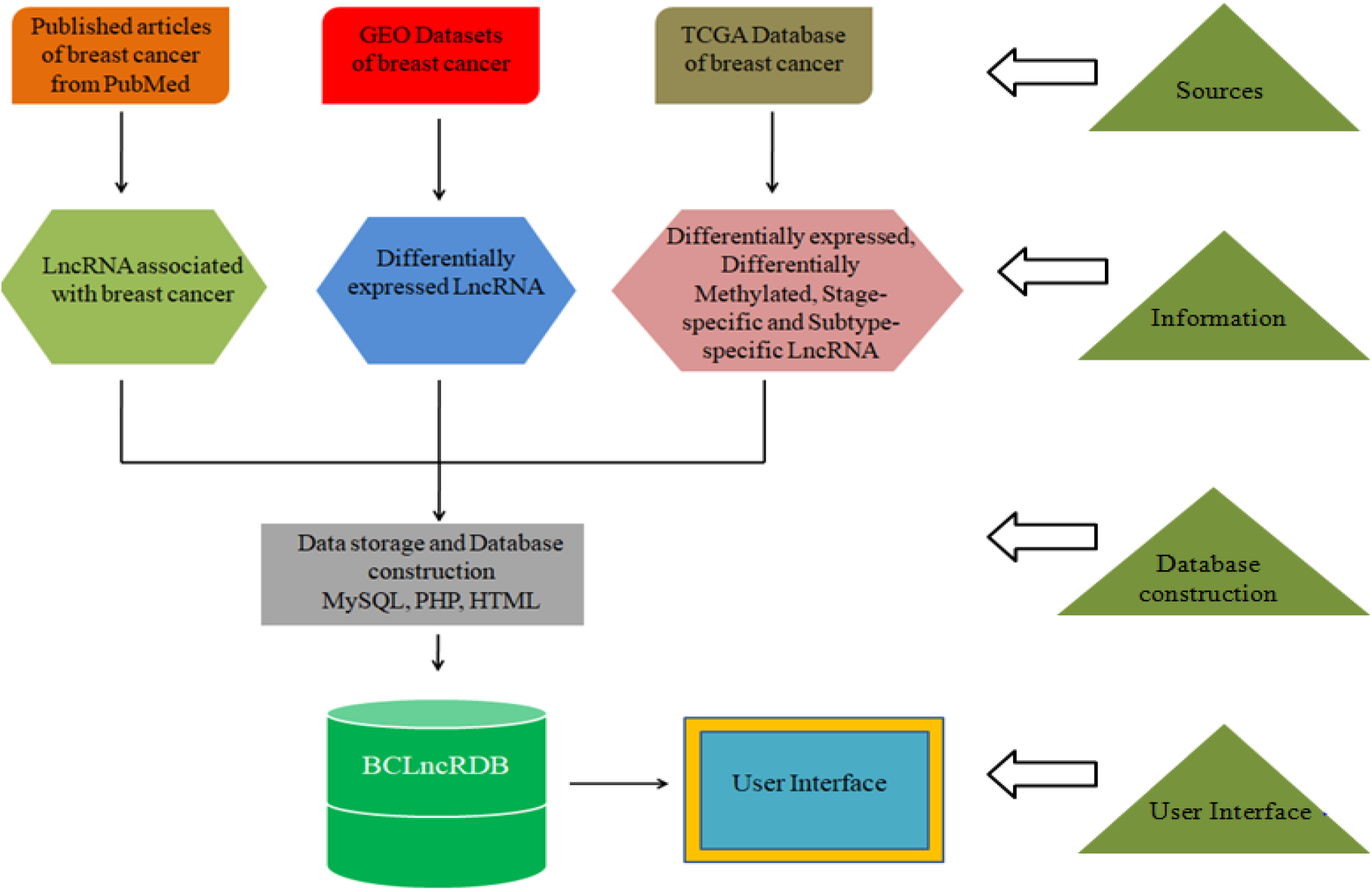
Schematic schema of database construction (BCLncRDB).

### Data collection from published literature

To collect valuable information on lncRNAs associated with breast cancer from various published literature, all literature (published till 14th May 2022) was extracted from the PubMed database using keywords such as ‘long non-coding RNA’, ‘lncRNA,’ and ‘long non-coding along with ‘breast.’ Secondly, all selected literature was curated manually, and filtered based on criteria such as (i) research articles containing experiments on breast cancer patient samples; (ii) samples were not treated with any chemicals or drugs; and (iii) experimental findings have been validated using expression data of breast cancer patients, cell-line data or animal models induced with breast cancer. All review articles and articles on experiments with samples treated with any chemicals or drugs were excluded from collecting information on lncRNAs associated with breast cancer.

### Data collection from GEO

To collect useful GEO Dataset, the same keywords were used to search GEO as used to search literature in PubMed. The GEO Datasets were retrieved based on the association of some criteria: (i) Organism should be Homo sapiens; (ii) Expression Profiling by array; (iii) Not treated with any drug/chemical; (iv) Tissue-specific. Those Datasets that did not match these criteria were discarded. Differentially expressed lncRNAs were identified by comparing expression levels of lncRNAs between tumor and normal tissues using the limma R package. Adjusted P-value < 0.05 and |log2fold-change| ≥ 2 were used to find significantly differentially expressed lncRNAs.

### Data collection from TCGA Database

Differential expression, differential methylation, subtype-specific, and stage-specific information of lncRNAs associated with breast cancer were extracted by analyzing various TCGA data. Differentially expressed lncRNA were identified by comparing expression levels of lncRNAs between tumor and normal tissues using the DESeq2 R package. Adjusted P-value < 0.05 and |log2fold-change| ≥ 2 were used to find significantly differentially expressed lncRNAs. Differentially methylated lncRNAs were identified by analyzing TCGA methylation data of breast cancer using the R/Bioconductor package ELMER [Silva et al., 2019].

### Other information on lncRNAs

Other information like Ensembl ID, sequences, and chromosome information of all lncRNAs associated with breast cancer were identified from various sources, including TCGA and Ensembl Database. Published literature from PubMed was used to retrieve information on the influence of drugs on lncRNA expression associated with breast cancer. The lncLocator database and published literature were used to find information on the subcellular localization of lncRNAs.

### Database Construction

All collected data on lncRNAs were stored and managed into a database named “BCLncRDB” using MySQL, a relational database structured query language; the user interface, or the front-end of the database, was built using HTML and PHP for browsing and searching the data contained therein.

## Acknowledgments

Vaibhav Vindal would like to acknowledge ICMR (ISRM/12(72)/2020, ID: 2020-2951), IoE UoH (No. UoH/IoE/RC3-21-052), and DBT (No. BUILDER-DBT-BT/INF/22/SP41176/2020) for their financial support. Swapnil Kumar would like to acknowledge ICMR for providing financial assistance as the Senior Research Fellowship (Grant No. 3/2/2/113/2019/NCD-III, ID: 2019-6723). Authors would like to thank Mr. Sayed Saleem Javed for his technical assistance in database hosting on the server.

## Data availability

The BCLncRDB is publicly available at http://sls.uohyd.ac.in/new/bclncrdb. The downloadable information of this database is available at http://sls.uohyd.ac.in/new/bclncrdb/Downloads.php.

